# Annulment of antagonism shifts properties that are beneficial to plants in two-member consortia of *Bacillus velezensis*

**DOI:** 10.1101/2021.10.13.464333

**Authors:** Jiahui Shao, Yan Liu, Jiyu Xie, Polonca Štefanič, Yu Lv, Ben Fan, Ines Mandic-Mulec, Ruifu Zhang, Zhihui Xu, Qirong Shen

## Abstract

*Bacillus* spp. strains that are beneficial to plants are widely used in commercial biofertilizers and biocontrol agents for sustainable agriculture. Generally, functional *Bacillus* strains are applied as single strain communities since the principles of synthetic microbial consortia constructed with *Bacillus* strains remain largely unclear. Here, we demonstrated that the mutual compatibility directly affects the survival and function of two-member consortia composed of *B. velezensis* SQR9 and FZB42 in the rhizosphere. A mutation in the global regulator Spo0A of SQR9 markedly reduced the boundary phenotype with wild-type FZB42, and the combined use of the SQR9∆*spo0A* mutant and FZB42 improved biofilm formation, root colonization and the production of secondary metabolites that are beneficial to plants. We further confirmed the correlation between the swarm discrimination phenotype between the two consortia members and the effects that are beneficial to plants in a greenhouse experiment. Our results provide evidence that social interactions among bacteria could be an influencing factor in achieving a desired community-level function.

**IMPORTANCE:** *Bacillus velezensis* is one of the most widely applied bacteria in biofertilizers in China and Europe. Additionally, the molecular mechanisms of plant growth promotion and disease suppression by representative model strains are well established, such as *B. velezensis* SQR9 and FZB42. However, it remains extremely challenging to design efficient consortia based on these model strains. Here, we showed that swarm discrimination phenotype is one of the major determinants affects the performance of two-member *Bacillus* consortia in vitro and in the rhizosphere. Deletion in global regulatory gene *spo0A* of SQR9 reduced the strength of boundary formation with FZB42, result in the improved plant growth promotion performance of dual consortium. This knowledge provides new insights into efficient probiotics consortia design in *Bacillus*.

## INTRODUCTION

Microorganisms do not exist alone in the rhizosphere and live surrounded by an enormous number of other organisms in close communities (1). Microbes are social creatures that interact with and coordinate the behaviors of each other, exhibiting various forms of relationships, including commensalism, neutralism, cooperation, competition, etc. (2). Social interactions among microbes can affect microbial community development and composition (3). Notably, bacterial interactions in the fly gut affect host fitness related to development, fecundity, and lifespan (4). Specifically, fly lifespan and gut microbiome abundances are markedly influenced by interactions between bacterial species (4). However, due to complexity of plant-associated microbiomes, the information on social interactions among bacteria in the rhizosphere and how they are linked to plant fitness has not been established yet.

For combination of probiotic bacterium, mutual compatibility is an essential trait required for their synergistic effects on host fitness, and cooperative behavior between individuals potentially enhanced their compatibility (5). Swarming is a bacterial cooperative behavior where billions of flagellated bacteria migrate together over solid surfaces (6). Studies have shown that cells of the soil bacteria *Bacillus subtilis* and *Myxococcus xanthus* can discriminate kin from nonkin in the context of swarming, where non-kin strains exhibit spatial segregation between swarms and kin merge (7, 8). Thus, it has been hypothesized that cooperative acts with a multitude of benefits are preferentially directed to highly related strains or kin (9). For *Bacillus*, three kinds of interaction phenotypes (merging, intermediate and boundary) could be observed on agar plates. Stefanic et al. (7) showed that cooperative behavior (merging of strains on swarming agar) is common among highly related strains of *B. subtilis* that share more than 99.5% identity at the *gyrA* genes and even between more distant *Bacillus* species (10). In contrast, antagonism/boundary prevails between *Bacillus* strains within species with less than 99.5% *gyrA* identity and between closely related species (10). This relatedness–sociality pattern has been also confirmed at the genome level of *B. subtilis* isolates (7). Additionally, it has been shown that *Bacillus* strains that merge their swarms also co-inhabit the plant root (7). Kin recognition or discrimination systems can elevate the kinship of relatives and thus stabilize cooperative behaviors (9).

Recent reports on kin discrimination (KD) system in *B. subtilis* suggested that this discrimination is achieved by nonkin exclusion rather than kin recognition (10, 11). *B. subtilis* cells produce a diverse array of toxins, and antibiotics form a blockade that only kin strains can survive. Moreover, several kin discrimination loci, including contact-dependent inhibition (CDI) protein (*wapAI*), cell-surface molecules (*lytC*, *dltA*, *tuaD*), and mobile elements such as phages, were identified by mutation (12). However, it is still unclear how these numerous factors work synergistically and contribute to kin discrimination. The blockade formed by close relatives prevents public goods from being exploited by nonkin cells (11, 13). In bacterial cooperation, public goods are generally defined as compounds that provide a collective benefit, usually through release to the extracellular environment (14). The production of public goods by *Bacillus*, such as extracellular matrix (EPS), siderophores and lipopeptide (e.g. surfactin), are controlled by a quorum sensing system (15, 16) and by global transcriptional regulators such as Spo0A and DegU (14, 17). Social interactions among *Bacillus* strains influence the production of public goods at the community level; however, to the best of our knowledge, few studies have focused on how antagonisms or cooperativity affect secretions of lipopeptide antibiotics and plant growth promoting hormones by bacteria in multicellular groups.

Among *Bacillus* spp., *B. velezensis* SQR9 and FZB42 are the most extensively studied strains for revealing beneficial plant-microbe interactions, including the stimulation of root growth, facilitation of nutrient uptake, and prevention of diseases in plants (18–20). In addition, both of these strains are used commercially as biofertilizers for agricultural production. In this work, we observed that the swarming phenotype between the well-established plant growth-promoting bacteria (PGPB) *B. velezensis* SQR9 and FZB42 switched from boundary to merging on agar plates after *spo0A* gene disruption. We further compared the properties that are beneficial to plants between mutual incompatibility consortia (SQR9 and FZB42) and mutual compatibility consortia (SQR9∆*spo0A* and FZB42). Our results demonstrated that microbial interactions affect community-level functions and behaviors in the rhizosphere.

## RESULTS

### *B. velezensis* SQR9 antagonism against FZB42 at the swarm encounter depends on Spo0A

Swarming motility is a cooperative movement involving the exchange of public goods (14, 15), which can serve as a model system to address social interactions between strains at the point of swarm encounter. The results showed that *B. velezensis* SQR9 swarm forms a prominent boundary line with *B. velezensis* FZB42 in the swarming assay (Fig. 1), which according to published work suggests that interactions between strains involves antagonism and that this two strains are non-kin (12, 21). We previously reported that a novel Sfp-dependent fatty acid, bacillunoic acid, produced by SQR9, directly inhibits the growth of FZB42 on agar plates (22). We therefore tested whether bacillunoic acid might be involved in the SQR9-FZB42 boundary formation. However, the ∆GI mutant of SQR9, which lacks the ability to biosynthesize bacillunoic acid, still exhibited a boundary phenotype similar to that of the wild-type strain (Fig. 1), which suggests that additional antagonistic factors contribute to SQR9-FZB42 boundary formation. Indeed, the SQR9∆*sfp* mutant showed slightly diminished boundary at the encounter with FZB42 compared to the wild-type strain (Fig. 1). Sfp is a 4’-phosphopantetheinyl transferase that has been proven to be essential for the production of most lipopeptides and polyketides in FZB42 (23). Therefore, results in figure. 1 suggest that Sfp-dependent secondary metabolites of SQR9 might contribute to the swarm boundary phenotype.

**Figure 1.**
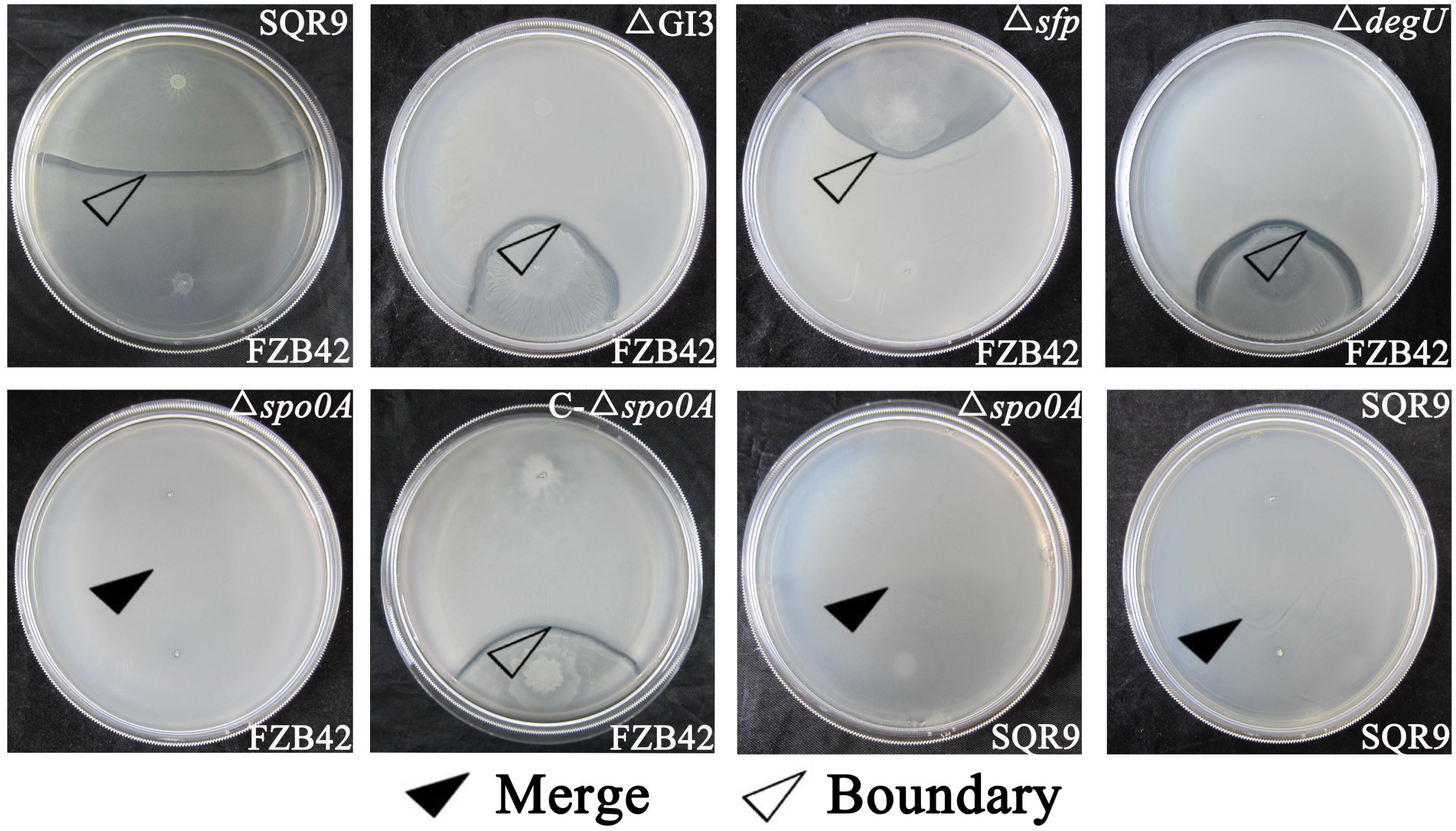
Different candidate gene deletions of *B. velezensis* SQR9 showed varied multicellular swarming phenotypes with wild-type *B. velezensis* FZB42. Results are representative of three experiments. For each photograph, *B. velezensis* SQR9 or its derivative strains were spotted on the surface of the swarming agar (B-medium) at similar distance appart, with either wild type *B. velezensis* FZB42 or SQR9 spotted on the lower portion of plates.

In *Bacillus* spp., the global regulators Spo0A and DegU jointly control the transcription of multicellular behavior such as biofilm formation (24), swarming and production of lipopeptide antibiotics (17, 24). Interestingly, deletion of *spo0A*, but not *degU*, shifted the swarming phenotype from boundary to merging in the swarming assay with *B. velezensis* FZB42 (Fig. 1). In contrast, deletion of *degU* preserved boundary. Moreover, the prominent boundary formed at increased distance from the point of SQR9 mutant inoculation to that of FZB42 (Fig. 1). Moreover, complementation of the *spo0A* gene in the SQR9∆*spo0A* mutant restored its ability to exhibit a boundary swarming phenotype with strain FZB42. Overall, we discovered that the regulator Spo0A in SQR9 is essential for boundary formation and potentially to its antagonism against FZB42.

### Antagonism between *B. velezensis* SQR9 and FZB42 affects biofilm formation and root colonization phenotypes in cocultures

A previous study demonstrated that kin strains formed mixed biofilms on roots, while nonkin strains engaged in antagonistic interactions, resulting in only one strain primarily colonizing the root (7). Although FZB42, SQR9 and SQR9∆*spo0A* mutant showed indistinguishable growth curves in monocultures (Fig. S1), we observed that coinoculation of SQR9 and FZB42 in MSgg liquid medium resulted in significant defects in biofilm formation (Fig. 2A, 2B & 2C) and strong antagonisms of SQR9 against FZB42, with only SQR9 remaining in coculture after 48 hours incubation (Fig. 2C). In contrast, the SQR9∆*spo0A* mutant, which formed thin and flat biofilms in monoculture regained in coculture with FZB42 a biofilm architecture indistinct from that of the monocultures (Fig. 2A). Moreover, the SQR9∆*spo0A* mutant and FZB42 coexisted in roughly equal proportion in coculture pellicle biofilms (Fig. 2B & 2C). This again suggests that deletion of *spo0A* alleviates the antagonisms of SQR9 against FZB42.

**Figure 2.**
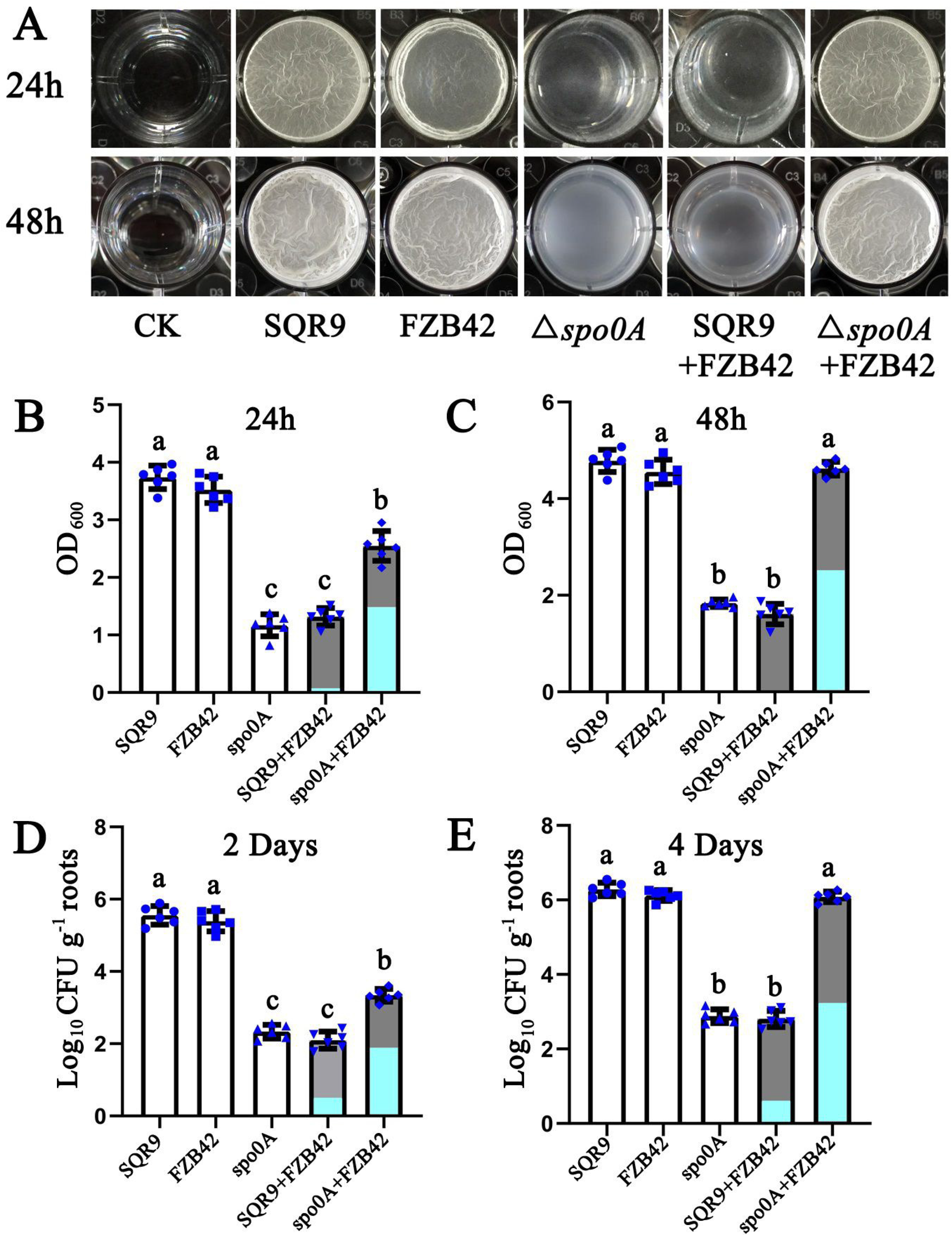
Pellicle biofilm formation and cucumber root colonization by monoculture and co-culture of *Bacillus* strains. SQR9 and FZB42 indicate wild type *B. velezensis* SQR9 and FZB42, respectively, and spo0A indicates the *spo0A* mutant of *B. velezensis* SQR9. (A) Floating pellicle formation by monoculture and co-culture of *Bacillus* strains. (B) and (C), OD_600_ of solubilized crystal violet from the microtiter plate assay at 24 and 48 hours, respectively. (D) and (E), the *Bacillus* populations of monoculture and co-culture colonizing cucumber seeding roots at 2 and 4 day time points. For co-culture treatments in B, C, D and E, the ratio of dark grey (SQR9 or *spo0A* mutant) and light blue (FZB42) indicate the proportion of different cells in the population, dark grey represents the cells of strain SQR9 or its *spo0A* mutant, light blue represents the cells of FZB42 wild type. Error bars indicate the standard deviations from the results from six independent experiments. Different letters above the bars indicate significant differences (*p*<0.05).

The root colonization pattern of these treatments correlated well with the measured values of pellicle biomass (Fig. 2D & 2E). After 2 days of incubation in the hydroponic system, the results showed that approximately 10^5^ CFU g^−1^ root of SQR9 or FZB42 cells were detected when inoculated as monocultures. However, only 10^2^ CFU g^−1^ root of SQR9∆*spo0A* was detected in monoculture and 10^2^ CFU g^−1^ root of the co-culture SQR9 with FZB42, with SQR9 again showing a fitness advantage over FZB42 (Fig. 2D). When SQR9∆*spo0A* and FZB42 were co-inoculated in the hydroponic system, cumulatively10^3^ CFU g^−1^ root was observed during the same period on the root, with slight competitive advantage of FZB42 over SQR9∆*spo0A* (Fig. 2D). After 4 days of incubation, root colonization by SQR9 and FZB42 reached a value of 10^6^ CFU g^−1^ root, while only 10^3^ CFU g^−1^ root of SQR9∆*spo0A* or wild type coculture (SQR9+FZB42 treatment) colonized the root with even more evident dominance of wild type SQR9 over FZB42 (Fig. 2E). However, the SQR9 dominance was lost when the SQR9∆*spo0A* mutant was used in coculture with FZB42, and lack of antagonism is also observed by high CFU counts of the coculture (10^6^ CFU g^−1^ root) (Fig. 2E). In summary, our results demonstrate that the swarming interaction phenotypes correlate with biofilm phenotype in liquid medium and on plant roots and are reflected in the community composition of a two-member *Bacillus* consortium.

### Avoidance of antagonism promotes synergism in production of secondary metabolites beneficial to plants

In *B. subtilis*, the kin discrimination system also influences antibiotic gene expression (12) and above results suggest that Spo0A regulated antibiotic synthesis, which may contribute to antagonistic social behaviour. We therefore tested whether Spo0A linked SQR9-FZB42 swarm phenotype involves lipopeptide production (bacillomycin D and fengycin), which is also a major weapon for the growth inhibition of pathogenic fungi in the rhizosphere (25, 26). To investigate this possibility, we first tested the antifungal activities of supernatants of monocultures and cocultures *in vitro* using the oxford cup-based agar diffusion assay. Both monocultures (SQR9 or FZB42) were more effective at antagonizing *Fusarium oxysporum* (FOC) than the coculture of SQR9 and FZB42. In line with the previous results on biofilm, the supernatant of SQR9∆*spo0A* mutant alone showed the lowest antifungal activity, in contrast the coculture of SQR9∆*spo0A* mutant and FZB42 showed the strongest antifungal activities visible as the clearing zone around the oxford cup (Fig. 3 & S2).

**Figure 3.**
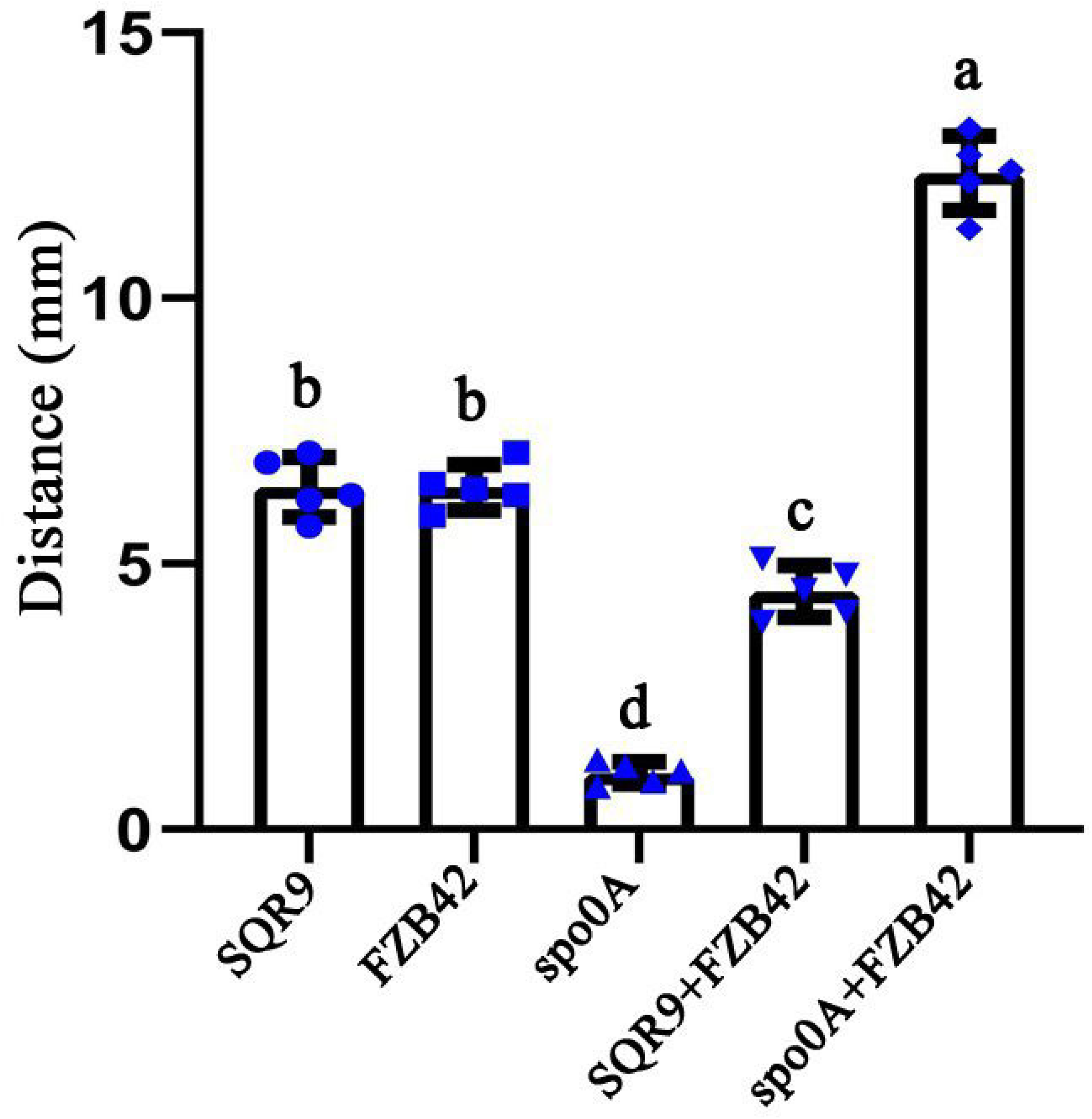
Distance between the fungal mycelium (*F. oxysporum*) and the wall of oxford cup filled with the supernatants of monoculture or co-culture of *Bacillus* strains. Error bars indicate the standard deviations from the results from six independent experiments. SQR9 and FZB42 indicate wild type *B. velezensis* SQR9 and FZB42, respectively, and spo0A indicates *spo0A* mutant of *B. velezensis* SQR9. Different letters above the bars indicate significant differences (*p*<0.05).

Previously, it was demonstrated that secretion of lipopeptides (bacillomycin D and fengycin) is the main mechanism by which SQR9 and FZB42 suppress the growth of *F. oxysporum* (27, 28). Additionally, methods for the detection of bacillomycin D and fengycin by RP-HPLC are well established (27). To further analyse the potential to exert antifungal activities *in vitro*, we monitored the production of bacillomycin D and fengycin in mono- and cocultures by HPLC. The results showed that the supernatants of the SQR9∆*spo0A* and FZB42 coculture showed the largest peak area of both bacillomycin D and fengycin, which can partly explain the strongest antifungal activities of this coculture *in vitro* (Fig. 4). Consistent with the above results the supernatant of the SQR9 and FZB42 coculture showed the smallest peak area of both bacillomycin D and fengycin (Fig. 4B).

**Figure 4.**
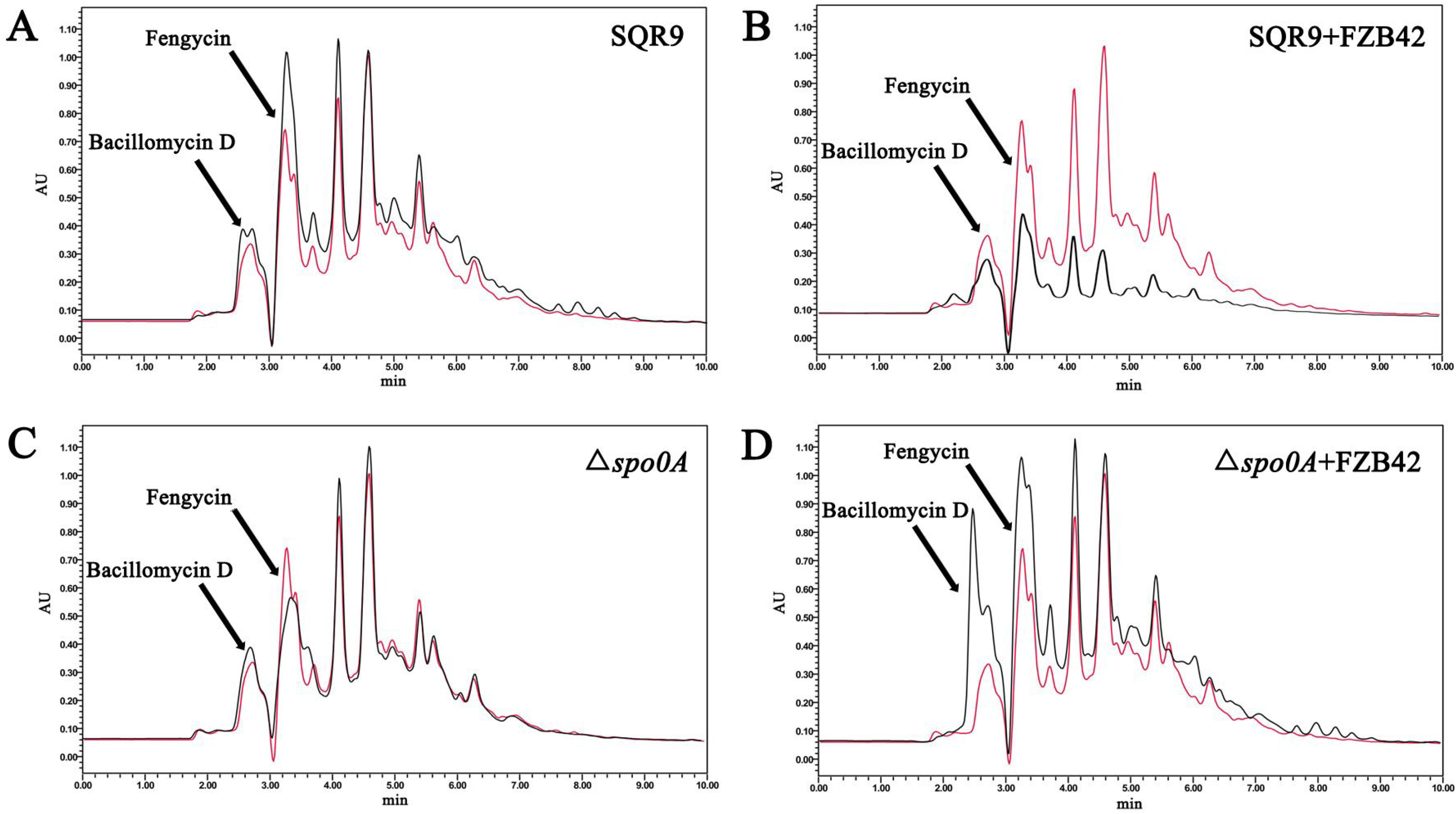
Reversed-phase HPLC chromatograms of lipopetides (bacillomycin D and fengycin) produced by monoculture and co-culture of *Bacillus* strains. For the better comparison, we used liquid chromatographic peaks of lipopetides produced by *B. velezensis* FZB42 as internal reference (red lines). Liquid chromatographic peaks in black indicate different samples: A, lipopetides produced by *B. velezensis* SQR9; B: lipopetides produced by co-culture of *B. velezensis* SQR9 and FZB42; C: lipopetides produced by *spo0A* mutant of *B. velezensis* SQR9; D: lipopetides produced by co-culture of *B. velezensis* SQR9∆*spo0A* and wild type FZB42.

The production of the phytohormones indole-3-acetic acid (IAA) and acetoin by *Bacillus* proved to be efficient in promoting plant growth (19), thus we tested whether the sociality of SQR9∆*spo0A*-FZB42 coculture also modulates plant hormone production. The coculture of SQR9∆*spo0A* and FZB42 showed the highest secretion of IAA and acetoin, while the coculture of SQR9 and FZB42 showed the lowest secretion at both metabolites at 24- and 48-hour time points (Fig. 5). In summary, the Spo0A mutant of SQR9, when in co-culture with the FZB42 positively affects the production of plant beneficial secondary metabolites by *Bacillus velezensis* strains, with similar patterns observed for IAA and acetoin production.

**Figure 5.**
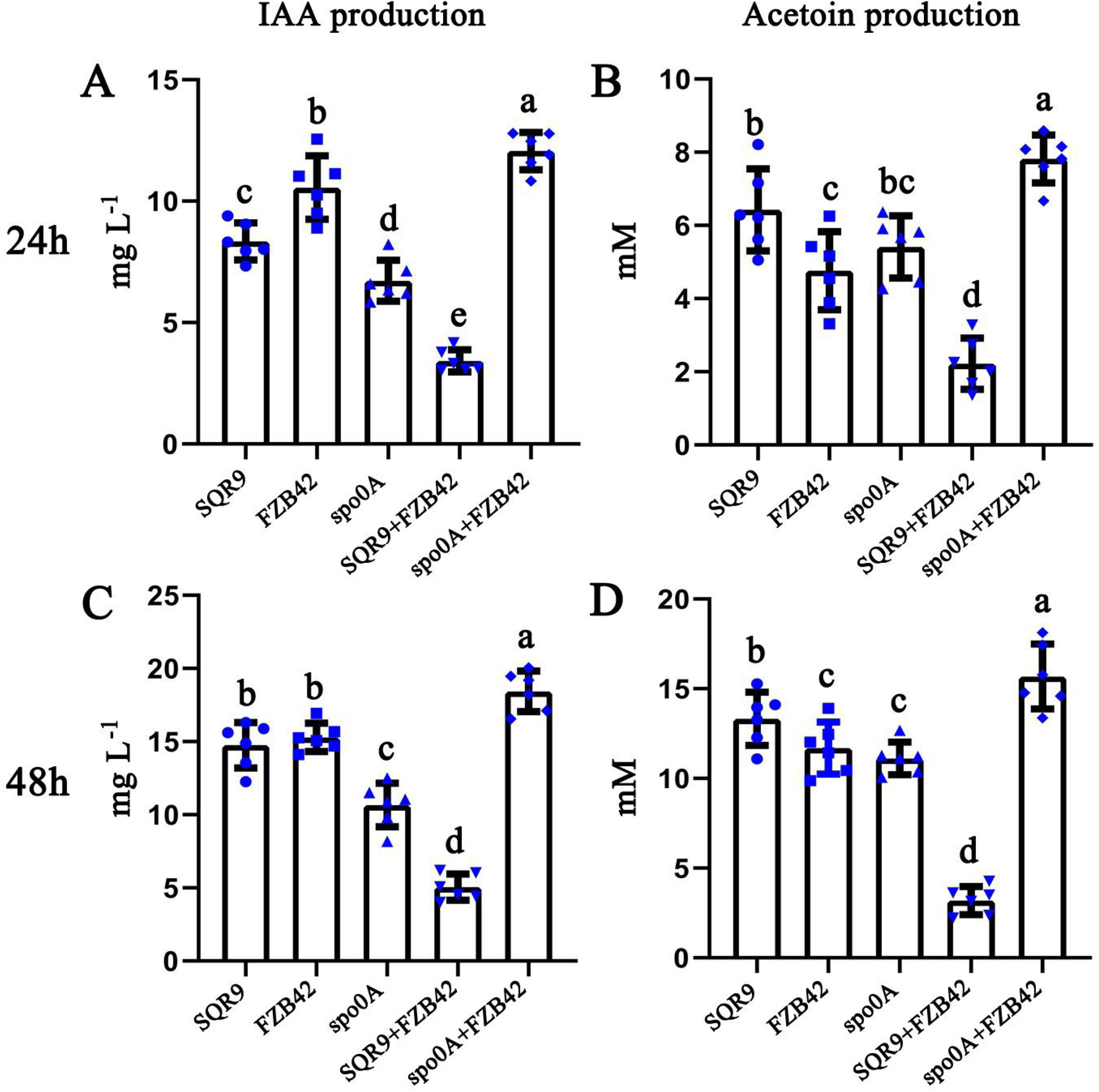
Production of IAA and acetoin in monoculture and co-culture of *Bacillus* strains. (A) and (C), IAA production of monoculture and co-culture of *Bacillus* strains was determined at 24 and 48 hours, respectively. (B) and (D), acetoin production in monoculture and co-culture of *Bacillus* strains at 24 and 48 hours’ time points, respectively. Error bars indicate the standard deviations from the results from six independent experiments. SQR9 and FZB42 indicate wild type *B. velezensis* SQR9 and FZB42, respectively, and spo0A indicates *spo0A* mutant of *B. velezensis* SQR9. Different letters above the bars indicate significant differences (*p*<0.05).

### Alleviation of antagonistic interactions of two-member *Bacillus* consortia influences its plant promoting properties

Root colonization and the production of secondary metabolites that are beneficial to plants are the two most important traits for efficient PGPB application (29, 30). We predicted that *Bacillus* consortia with mutual compatibility will perform better as biocontrol agents against pathogens than incompatible *Bacillus* consortia. The addition of monocultures and cocultures of SQR9 and FZB42 improved the plant height and shoot dry weight to varying degrees compared with cucumber plants inoculated with FOC (Fig. 6). The plant shoot height and shoot dry weight analysis showed that the coculture of SQR9∆*spo0A* and FZB42 performed the best (Fig. 6B & 6C), the monoculture of SQR9 was second best (Fig. 6B & 6C), and other treatments (FZB42, SQR9∆*spo0A*, SQR9+FZB42) ranked third in terms of plant growth promoting effects (Fig. 6B & 6C). Overall, plants inoculated by SQR9∆*spo0A* mutant and FZB42 mixed consortia showed the best growth under the FOC pathogen stress as both the plant height and shoot dry weight were almost identical to those of cucumber plants without the addition of FOC to the soil (Fig. 6).

**Figure 6.**
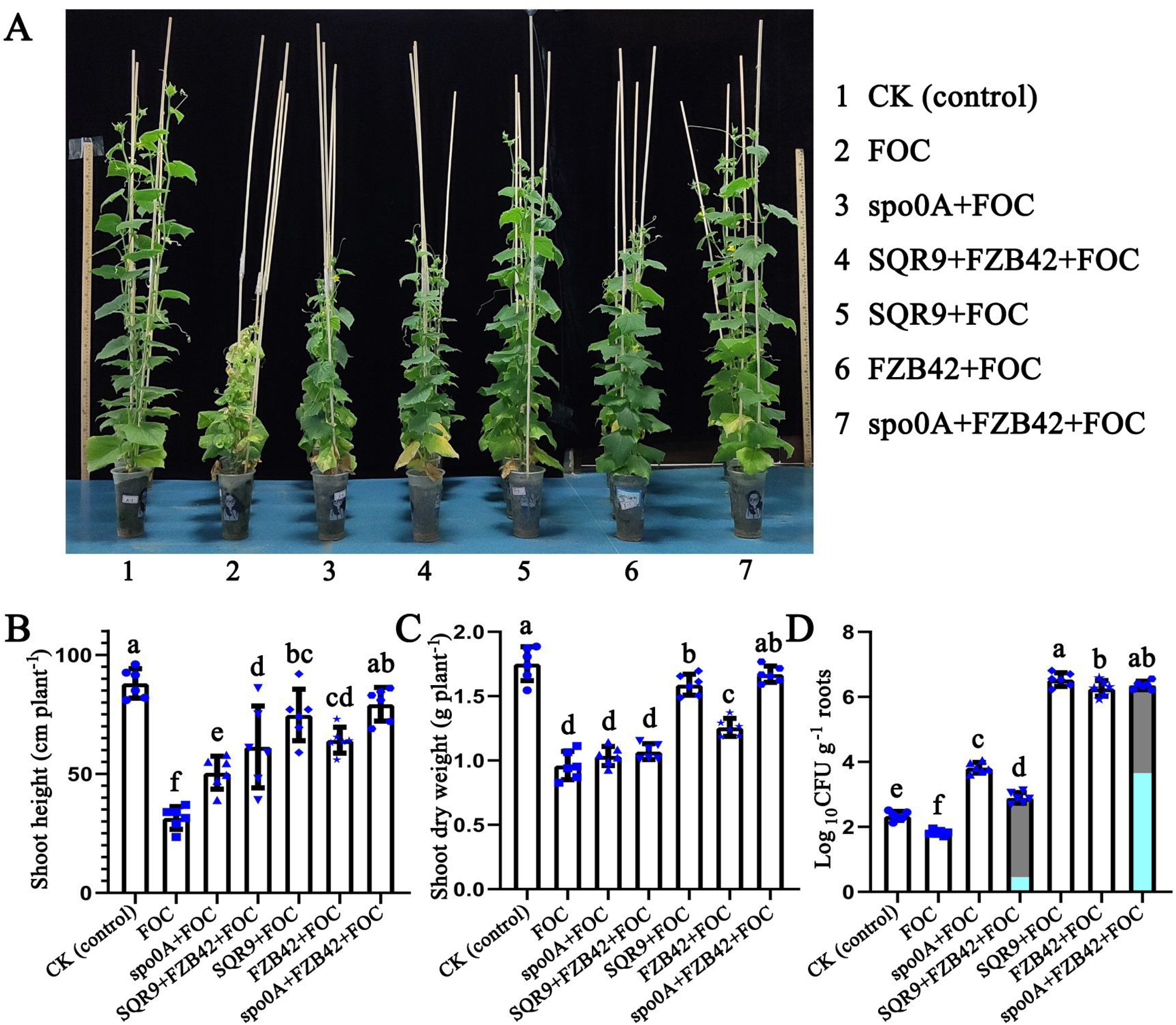
Greenhouse experiment indicated that combined used of SQR9∆*spo0A* and FZB42 significantly improved the growth of cucumber under FOC pathogen (*F. oxysporum*) stress. SQR9 and FZB42 indicate wild type *B. velezensis* SQR9 and FZB42, respectively, and spo0A indicates *spo0A* mutant of *B. velezensis* SQR9. (A) Representative images of the cucumber plants inoculated with monoculture and co-culture of *Bacillus* strains before harvest. (B) and (C) show the effect of different treatments on cucumber plant height and shoot dry weight, respectively. (D) *Bacillus* populations of monoculture and co-culture colonizing cucumber roots after harvest. For co-culture treatments (SQR9+FZB42+FOC and spo0A+FZB42+FOC), the ratio of dark grey (SQR9 or *spo0A* mutant) and light blue (FZB42) indicate the proportion of different cells in the population, dark grey represents the cells of SQR9 or its *spo0A* mutant, light blue represents the cells of FZB42. Error bars indicate the standard deviations from the results from six independent experiments. Different letters above the bars indicate significant differences (*p*<0.05).

To further investigate the mechanism underlying the plant promotion effect, we monitored the number of bacterial cells in the rhizosphere soil before harvest (50 days old plants). The results showed that both strains SQR9 and FZB42 could be detected in the coculture (SQR9+FZB42) treatment; however, the total number of cells was significantly lower than that of SQR9 or FZB42 monocultures (Fig. 6D). Cell number detected in coculture (SQR9∆*spo0A*+FZB42) treatment compared to monoculture of SQR9 or FZB42, and roughly equal amounts of strain SQR9∆*spo0A* and FZB42 coexisted in the rhizosphere community (Fig. 6D) while in SQR9+FZB42+FOC treatment, the strain SQR9 dominated the community colonizing the root (Fig. 6D). Overall, greenhouse experiments confirmed that the social interactions (swarming incompatibilities) indeed affected the activities of two-member *Bacillus* consortia that are beneficial to plants.

## DISCUSSION

The application of synthetic microbial communities is a novel trend for developing robust and stable microbial fertilizers (31). However, less is known about how to manipulate and improve the function of the designed microbial communities. Mutual compatibility could be one of fundamental principles for rationally designed microbial consortia. Bacterial social interactions, such as antagonism and cooperation, are ubiquitous in microbial communities, and cooperation is known to facilitate the maintenance (32). Here, we demonstrated that antagonism between strains in the swarming assay influences cooperative and function of two-member consortia in the cucumber rhizosphere. The combined application of *Bacillus* strains whose swarms form a clearly visible boundary on swarming agar decreased the total abundance of *Bacillus* in the rhizosphere and reduced the production of bioactive secondary metabolites, resulting in decreased benefits for plants. Astonishingly, diminished boundary line between *Bacillus* swarms correlated with the beneficial effects of SQR9∆*spoOA*-FZB42 consortia on FOC infected plants and the consortia dependent FOC was as efficient as SQR9 monoculture treatment. Moreover, SQR9∆*spo0A* or FZB42 monocultures did not show comparable protection against FOC. Understanding the role of social interactions in community-level function is thus important for synthetic microbial community design.

For cooperation behaviours, bacteria usually exhibit KD-like behavior, in particular while engaged in swarming (10). The KD system in environmental bacteria is usually correlated with the production of bacteriocins (33), and the cost trade-off between bacteriocins and the secretion of secondary metabolites that are beneficial to plants could cause the poor effects of this approach on benefits for plants. Indeed, our results showed that the production of plant beneficial secondary metabolites (bacillomycin D, fengycin, IAA and acetoin) are highly correlated with the swarm discrimination phenotype in a two-member consortia of *Bacillus velezensis*. This idea is also supported by the observation that co-inoculation of antagonistic non-kin *B. subtilis* strains leads to strain exclusion on plant roots (7). Our results are consistent with this observation and show partial exclusion of FZB42 by SQR9 and coexistence of SQR9∆*spo0A* mutant with SQR9. We therefore suggest that combined application of antagonistic PGPR strains could have negative effects on its plant beneficial function realization and survival.

An important feature of the kinship dependent cooperation is sharing of ‘public goods’ that benefit all cells irrespective of which cells produce them (10). A loss of antagonisms and the concomitant increase in the production of secondary metabolites that are beneficial to plants (IAA, acetoin, lipopeptides) in coculture (SQR9∆*spo0A*+FZB42) may in part explain the beneficial effect on plants. It has been reported that the amphipathic lipopeptide surfactin acts a ‘public good’ produced by *B. subtilis* and that it is necessary for multicellular swarming by reducing the water surface tension (16). In this study, this function may be supplemented by the lipopeptides bacillomycin D and fengycin, which can also act as biosurfactants and could be therefore potential ‘public goods’ in coculture environments (34). Most importantly, previous studies demonstrated that bacillomycin D and fengycin are the major compounds that inhibit the growth of FOC by both SQR9 and FZB42 (27, 28). For plant growth-promoting (PGP) compounds, due to the limited knowledge of how biosynthesis of IAA and acetoin are regulated in *Bacillus*, it is difficult to disentangle the correlation between the KD like behavior and PGP compound production, but our results are in line with the prediction that *spo0A* potentially controls synthesis of both plant hormones.

Here, we observed that SQR9 wild type formed boundary but SQR9∆*spo0A* mutant merged with the wild-type FZB42 (Fig. 1). According to recent reports boundary formation is tightly associated with antagonism (14, 21) at the swarm encounter. Our results also suggest that SQR9 and FZB42 engage in antagonisms and that lack of Spo0A in SQR9 reduces the efficiency of the antimicrobial attack and defense systems. Although further investigations are required to identify the underlying molecular mechanism by which *spo0A* is involved in the KD like response during swarming, it is known that Spo0A is the master regulator that is dependent on the phosphorylation state (35) and that phosphorylated Spo0A is needed to induce sporulation, synthesis of several antibiotics and production of extracellular matrix (36). In *B. subtilis*, the lack of extracellular matrix impacted kin dependent recognition with the parental strain (14), but the absence of the matrix polysacharide in the mutant strain resulted in a boundary phenotype when competed against the parental strain. This differs from our results where the *spo0A* mutant, known to produce less matrix components (24) still merged with the parental strain and even with the non-kin. It seems that disappearance of a boundary between SQR9∆*spo0A* mutant and FZB42 removed the SQR9 dependent antagonism during the swarm encounter and simultaneously promoted the production of plant beneficial antifungals and hormones.

Besides Spo0A the DegS/U two-component system acts as a global regulatory system in *Bacillus subtilis*, where phosphorylated DegU is involved in regulation of genetic competence, swarming motility, biofilm formation and exoprotease secretion, and a small protein DegQ modulates DegU phosphorylation (37, 38). However, in *Bacillus velezensis* SQR9, according to our results *degU* mutant still displayed swarming motility and formed stronger boundaries with FZB42 on swarming agar. One possibility is that deletion of the *degU* gene improved the production of an antibiotic exerting its activity at the swarm encounter. Previous studies discovered that disruption of *degQ* gene in *Bacillus subtilis* NCD-2 increased the production of fengycin and surfactin (39) but whether these two antibiotics and DegU/S system contribute to boundary formation between non-kin strains remains to be studied in the future.

In conclusion, our findings suggest that the swarm discrimination phenotype between PGPR strains may reflect the community level effects when used as consortia for plant protection. Although our observation is only based on interactions between two plant beneficial strains, SQR9 and FZB42, they suggest that strains that antagonize at the swarm encounter also engage in competition on plant roots which may diminish their beneficial effect on plants. To prove this hypothesis more strains need to be tested, but our results imply that swarm interaction may serve as a predictive read out for synergistic effects between inoculants, which may have important implications for the design and application of synthetic *Bacillus* communities in concrete applications.

## MATERIALS AND METHODS

### Strains and growth conditions

The strains used in this study are listed in Table 1. *B. velezensis* strain SQR9 (CGMCC accession no. 5808; China General Microbiology Culture Collection Center) was used throughout this study. *B. velezensis* FZB42 and green fluorescent protein (GFP)-labeled *B. velezensis* FZB42 were acquired as kind gifts from Ben Fan (Nanjing Forestry University) and through the *Bacillus* Genetic Stock Center. *B. velezensis* strains and their mutants were grown in Luria-Bertani (LB) medium. Biofilm assays were performed in 24-well plates with 2 mL of MSgg medium (5 mM potassium phosphate, 100 mM morpholinepropanesulfonic acid (MOPS) pH 7, 2 mM MgCl_2_, 700 μM CaCl_2_, 50 μM MnCl_2_, 50 μM FeCl_3_, 1 μM ZnCl_2_, 2 mM thiamine, 0.5% glycerol, 0.5% glutamate, 50 μg mL^−1^ tryptophan, 50 μg mL^−1^ phenylalanine, and 50 μg mL^−1^ threonine) (40). For lipopeptide production and HPLC characterization, *B. velezensis* strains were grown in Landy medium (20 g L^−1^ glucose, 5 g L^−1^ L-glutamic acid, 1 g L^−1^ KH_2_PO_4_, 1 g L^−1^ yeast extract, 0.5 g L^−1^ MgSO_4_ 7H_2_O, 0.5 g L^−1^ KCl, 5 mg L^−1^ MnSO_4_ H_2_O, 0.16 mg L^−1^ CuSO_4_ 7H_2_O, 0.15 mg L^−1^ FeSO_4_ 7H_2_O, 2 mg L^−1^ L-phenylalanine, 1 g L^−1^ L-tryptophan, pH 7.0) (41). Antibiotics were added as required at the following concentrations: 20 μg mL^−1^ zeocin, 5 μg mL^−1^ chloramphenicol (Cm), 5 μg mL^−1^ erythromycin (Em) and 100 μg mL^−1^ ampicillin.

**Table 1.**
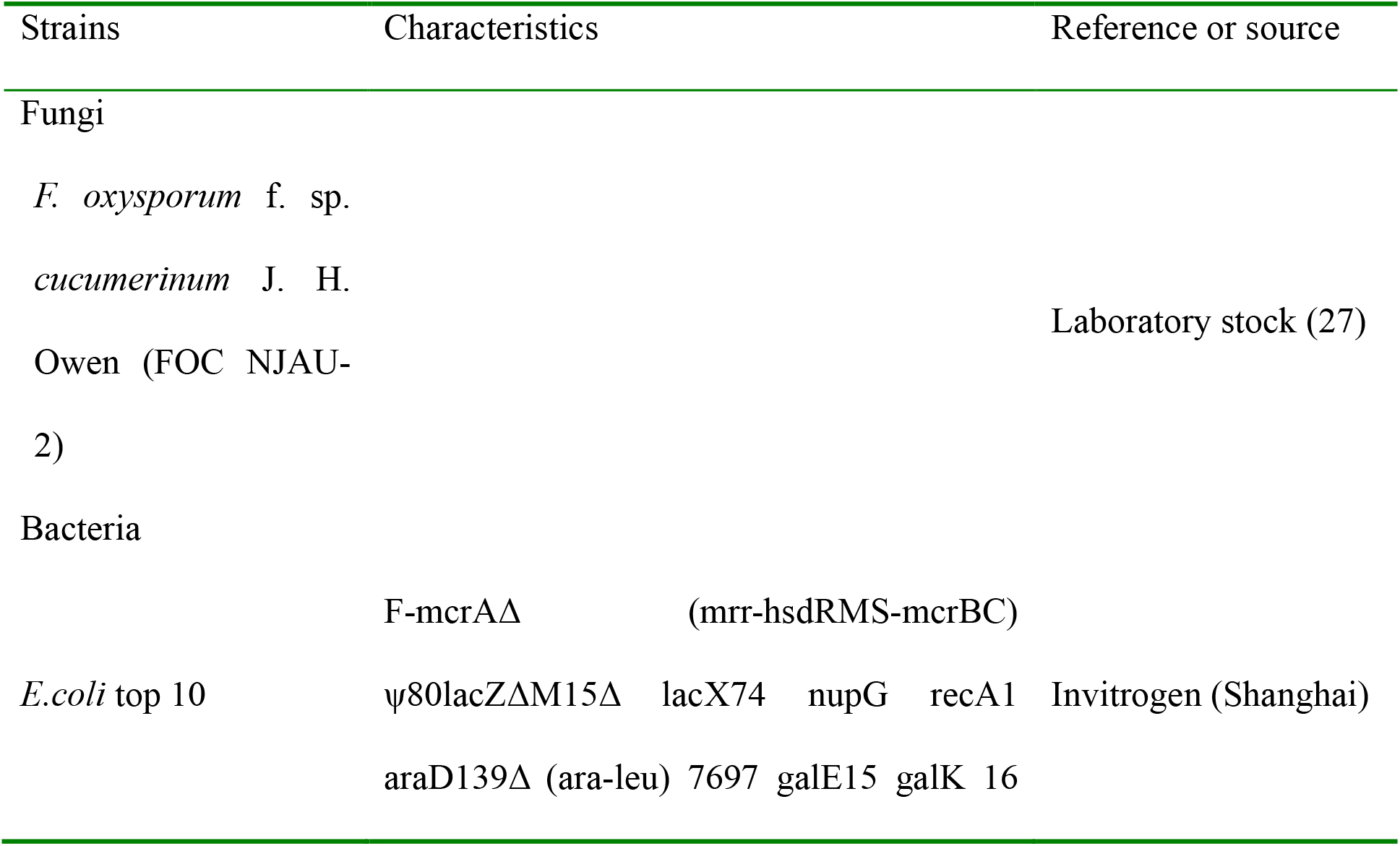
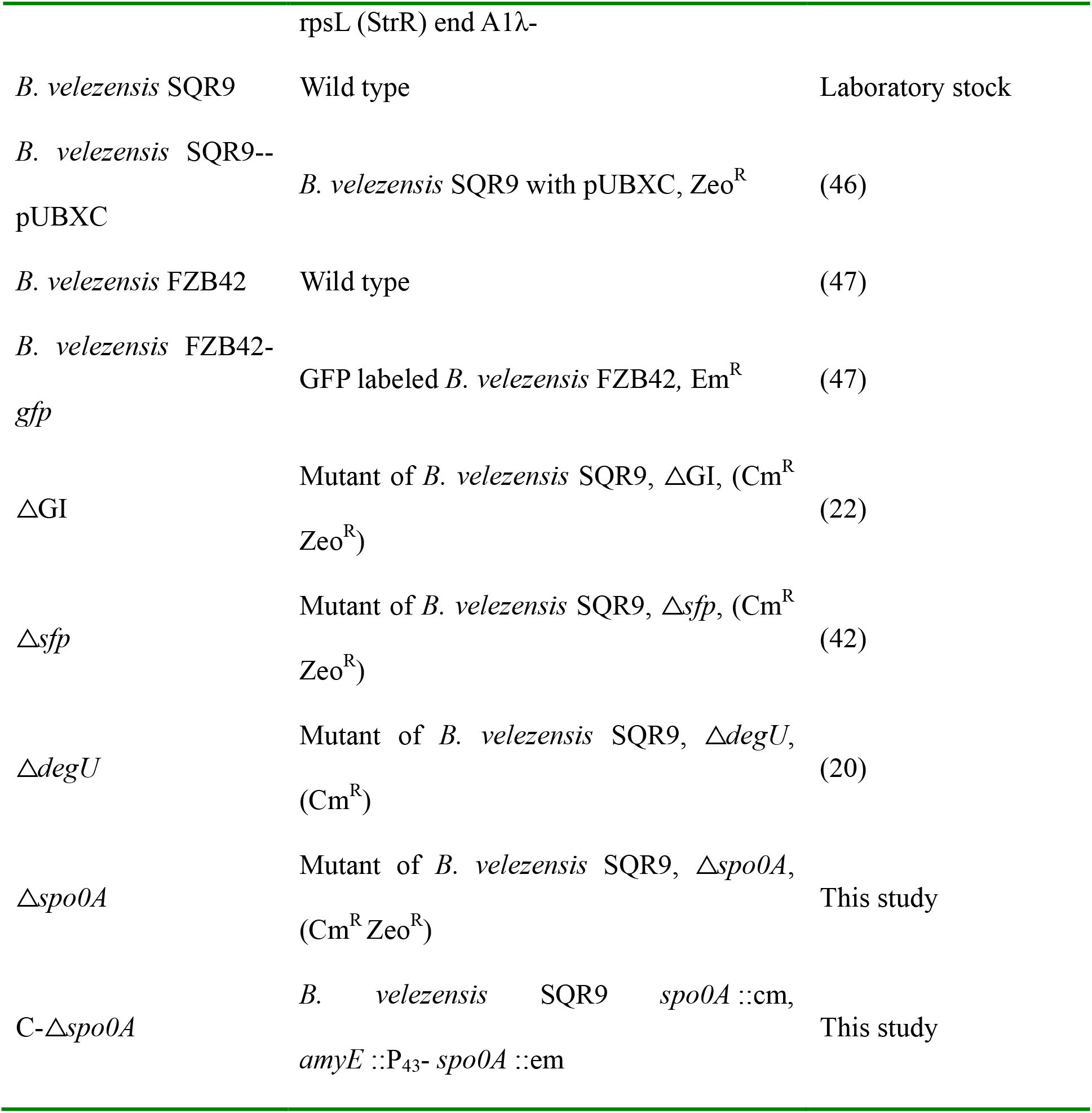
Microorganisms used in this study

### Swarming boundary assay

To test the social interaction between approaching swarms of wild-type *B. velezensis* FZB42, SQR9 and its mutants, 9-cm plates containing B-medium with 0.7% agar were freshly prepared (7). Strains were grown on solid Luria-Bertani (LB) plates at 30 °C for 16 h before use and then transferred to 3 mL of liquid B-medium and shaken overnight at 30 °C. The overnight cultures were then diluted to an optical density (OD_600_) of 0.5, and 2 μL was spotted on the plates at eat each side of the agar plate. The plates were dried in a laminar flow hood for 20 min, incubated for 2 days at 30 °C, and photographed. Three phenotypes (merge, intermediate and boundary) were assigned from the photos (7).

### Growth curve measurement

Growth curve experiments were assessed in 200 μL minimal medium-glucose-yeast (MSgg) medium in 96-well microtiter plates. The initial inoculum size was set at an OD_600_ value of 0.05. The OD_600_ was measured every 30 min at 30 °C with a Bioscreen C Automated Microbiology Growth Curve Analysis System (Growth Curve, USA). This assay was repeated three times.

### Construction of the marker-free deletion of *spo0A* gene in SQR9

The deletion of the *spo0A* gene was constructed using the *Pbc-pheS^#^-cat* (PC) cassette and overlap-PCR-based strategy as previously described (42). The construction of the PC cassette was carried out as described by Zhou et al. (43). Briefly, fragments including homologous sequences and PC cassettes were constructed by using overlap-PCR and then directly transformed into strain SQR9. The transformants were selected on LB plates containing Cm (5 μg/mL). The chloramphenicol-resistant colonies were cultivated to an OD_600_ of 1.0 without Cm, and a 100 mL aliquot of a 10-fold dilution of cultures (approximately 10^5^ cells) was plated on MGY-Cl medium (42). Targeted mutants were further confirmed by DNA sequencing.

### Biofilm assays

The pellicle biofilm assay was carried out in 24-well microtiter plates insert with 100 μm Sterile Nylon Mesh Cell Strainers (Biologix Cat#15-1100) in MSgg liquid medium, as described by Hamon and Lazazzera (44). After incubation, the cell strainer was taken out, pellicle biofilm formation was quantified by staining with crystal violet (CV). Cells in the pellicle biofilm were stained with CV, and then, unbound CV was removed with distilled water. The remaining CV was solubilized with 80% ethanol-20% acetone (1 mL). The absorbance of CV was measured using the SpectraMax i3x analysis system (Molecular Devices Corporation, CA) at 570 nm. For quantification of SQR9 and FZB42 cells in the pellicle biofilm, we conducted this experiment by labeled strains: using SQR9-pUBXC (carrying zeocin resistance gene) and GFP-labeled FZB42 (carrying erythromycin resistance gene). All cells in the pellicle which attached to the cell strainer were analyzed using the plate counting method with corresponding antibiotics.

### Bioassay of antagonistic activities against *F. oxysporum* of supernatants of monocultures and cocultures

*B. velezensis* cells were grown in Landy medium at 30 °C for 48 h, and LPs were then isolated by acid precipitation with concentrated HCl at pH 2.0. The precipitate was recovered by centrifugation at 8,000 ×g for 20 min, washed twice with acidic deionized water (pH 2.0), and then extracted twice with methanol (27). The pooled extraction solution was filtered through a 0.22 μm pore hydrophilic membrane, and a volume of 50 μL of extraction solution was dropped into an Oxford cup placed 2 cm from the edge of a petri plate and allowed to diffuse into agar. A plug (about 1 cm in diameter) of *F. oxysporum* from a 5-day-old (25°C) PDA plate of the growing *F. oxysporum* was placed in the center. The plates were then incubated at 25°C, and the distance between the edges of the petri dish and the fungal mycelium were measured after 5 days. The same volume of methanol was used as control. The experiment was repeated three times.

### Detection of lipopeptides (LPs) by reverse-phase (RP) HPLC

To compare the production of LPs produced by monocultures and cocultures of *Bacillus* strains, the supernatant was analysed by reversed-phase high-performance liquid chromatography (RP-HPLC) in a previous study (20, 27). Briefly, cells were grown in Landy medium at 30 °C for 48 h, and LPs were isolated as described above. The pooled extraction solution was filtered through a 0.22 μm pore hydrophilic membrane and then dried with a rotary vacuum evaporator. Finally, the residue was dissolved in 1 mL of phosphorous buffer (PBS; 0.01 M [pH 7.4]).

The determination conditions of bacillomycin D and fengycin were established by injecting 10 μL samples into an HPLC column (Eclipse XDB-C18, 5 μm; Agilent, Santa Clara, CA). The temperature was maintained at 30 °C during the experiment. The run was performed with a flow rate of 0.75 mL/min and a gradient of solvent A (0.1% [vol/vol] HCOOH) and B (CH_3_CN) and then 100% B after 20 min. To equilibrate the column, it was treated with 5% CH_3_CN-HCOOH for 3 min. A UV detector was used to detect the target peaks at 230 nm (27).

### IAA production

We grew individual *Bacillus* cells and their consortia in liquid Landy medium for 72 h at 22 °C in the dark with constant shaking (90 rpm). Bacterial cultures were then centrifuged (at 10000 g for 5 min), and the IAA concentration of the supernatants (ng/mL) was measured with an IAA ELISA Kit (R&D, Shanghai, China) following the manufacturer’s protocol (19). This assay was repeated three times.

### Acetoin production

Detection of acetoin in the monocultures and cocultures of *Bacillus* strains were performed by the method of Nicholson (45) as follows: one hundred forty microliters of creatine (0.5% [w/v] in water), 200 μL of α-naphthol (5% [w/v] in 95% ethanol), and 200 μL of KOH (40% [w/v] in water) were sequentially added to 200 μL of acetoin standard solution or appropriately diluted culture supernatant. The mixed samples were vortexed after each addition. The reaction mixtures were vortexed again after incubation for 15 min at room temperature before the measurement of A_560_.

### Root colonization

Each *Bacillus* strain and consortia of two strains were inoculated into the culture of two-week old sterile cucumber seedlings grown in 1/4 Murashige and Skoog (MS) culture medium. After 2 and 4 days, the *Bacillus* cells that had colonized the cucumber roots were collected and quantified using the method described by Xu et al. (42). The number of attached cells (SQR9-pUBXC and GFP-labelled FZB42) were analysed using the plate counting method with media supplemented with corresponding antibiotics.

### Greenhouse experiment

The greenhouse experiment was conducted from 25 July to 20 September 2020 at Nanjing Agricultural University. The soils used for the pot experiments were collected from a field with a history of cucumber cultivation with the following properties: pH 5.9; organic matter, 25.3 g kg^−1^; available N, 166.5 mg kg^−1^; available P 127.8 mg kg^−1^; available K, 256.8 mg kg^−1^; total N, 1.9 g kg^−1^; total P, 1.7 g kg^−1^; and total K, 15.2 g kg^−1^. The surfaces of cucumber seeds (Jinchun No. 4) were disinfected in 2% sodium hypochlorite for 4 min and then germinated in seedling trays at 28 °C. Two-week-old seedlings were transplanted into pots with 600 g of sterilized soil. Seven treatments were designed as follows: (1) CK (control, sterilized soil without inoculation); (2) FOC (sterilized soil inoculated with *F. oxysporum*); (3) SQR9+FOC (sterilized soil first inoculated with *F. oxysporum* then with *B. velezensis* SQR9); (4) FZB42+FOC (sterilized soil first inoculated with *F. oxysporum* then with *B. velezensis* FZB42); (5) Spo0A+FOC (sterilized soil first inoculated with *F. oxysporum* then with ∆*spo0A* mutant of SQR9); (6) SQR9+FZB42+FOC (sterilized soil first inoculated with *F. oxysporum* then with *B. velezensis* SQR9 and FZB42, respectively); and (7) Spo0A+FZB42+FOC (sterilized soil first inoculated with *F. oxysporum* then with *B. velezensis* ∆*spo0A* mutant of SQR9 and wild type FZB42, respectively). All strains were individually mixed into the soil as follows. Two weeks before transplant, *F. oxysporum* was first inoculated into the soil at 10^5^ spores g^−1^ soil. One week after transplantation, *B. velezensis* cells (SQR9, FZB42 and ∆*spo0A*) were inoculated into soil at 10^7^ cfu g^−1^. For the SQR9+FZB42 and Spo0A+FZB42 treatments, the SQR9 or SQR9∆*spo0A* mutant was inoculated first, the strain FZB42 was inoculated 2 days later, and both of them reached a cell density of 10^7^ cfu g^−1^ of soil. Each treatment was replicated 6 times. The cucumber plant was incubated in a growth chamber at 30 °C under a 16 h light regimen and irrigated with 1/4 h Hoagland medium.

### Statistical analysis

Differences among the treatments were calculated and statistically analysed with a one-way analysis of variance (ANOVA). Duncan’s multiple-range test was used when the one-way ANOVA indicated a significant difference (*p*<0.05). All statistical analyses were performed with IBM SPSS Statistics 20.

## ACKNOWLEDGEMENTS

This work was financially supported by the National Nature Science Foundation of China (31972512, 32072675 and 32072665), the Fundamental Research Funds for the Central Universities (KYXK202009). IMM and PS were supported by the Slovenian Research program P4-0116 and the projects J4-9302 and J4-8228.

## Data Accessibility

The accession numbers of the genome sequence of *Bacillus amyloliquefaciens* SQR9 and FZB42 in the NCBI are: CP006890.1 and CP000560.2.

## Author contributions

JS, YL, ZX designed the study, and JS, YL, JX, YL performed the experiments. JS, YL, ZX, JX analyzed the data and created the figures. JS and YL wrote the first draft of the manuscript, and ZX, IMM, PS, BF, RZ and QS revised the manuscript.

---

Figure S1. Growth curves of *B. velezensis* FZB42, *B. velezensis* SQR9 and its mutants strains in liquid MSgg medium.

Figure S2. Representative images of distance between the fungal mycelium (*F. oxysporum*) and the wall of oxford cup filled with the supernatants of monoculture or co-culture of *Bacillus* strains. SQR9 and FZB42 indicate wild type *B. velezensis* SQR9 and FZB42, respectively, and spo0A indicates *spo0A* mutant of *B. velezensis* SQR9. The solvent control is methanol.

## REFERENCES

1. Wall D. 2016. Kin recognition in bacteria. Annu Rev Microbiol 70:143–160. 10.1146/annurev-micro-102215-095325.

2. West SA, Diggle SP, Buckling A, Gardner A, Griffin AS. 2007. The social lives of microbes. Annu Rev Ecol Evol Syst 38:53–77. 10.1146/annurev.ecolsys.38.091206.095740.

3. Kong W, Meldgin DR, Collins JJ, Lu T. 2018. Designing microbial consortia with defined social interactions. Nat Chem Biol 14:821–829. 10.1038/s41589-018-0091-7.

4. Gould AL, Zhang V, Lamberti L, Jones EW, Obadia B, Korasidis N, Gavryushkin A, Carlson JM, Beerenwinkel N, Ludington WB. 2018. Microbiome interactions shape host fitness. Proc Natl Acad Sci U S A 115:E11951–E11960. 10.1073/pnas.1809349115.

5. Palmieri D, Vitullo D, De Curtis F, Lima G. 2017. A microbial consortium in the rhizosphere as a new biocontrol approach against *Fusarium decline* of chickpea. Plant Soil 412:425–439. 10.1007/s11104-016-3080-1.

6. Yan J, Monaco TH, Xavier JB. 2019. The ultimate guide to bacterial swarming: An experimental model to study the evolution of cooperative behavior. Annu Rev Microbiol 73:293–312. 10.1146/annurev-micro-020518-120033.

7. Stefanic P, Kraigher B, Lyons NA, Kolter R, Mandic-Mulec I. 2015. Kin discrimination between sympatric *Bacillus subtilis* isolates. Proc Natl Acad Sci U S A 112:14042–14047. 10.1073/pnas.1512671112.

8. Wielgoss S, Fiegna F, Rendueles O, Yu YTN, Velicer GJ. 2018. Kin discrimination and outer membrane exchange in *Myxococcus xanthus*: A comparative analysis among natural isolates. Mol Ecol 27:3146–3158. 10.1111/mec.14773.

9. Strassmann JE, Gilbert OM, Queller DC. 2011. Kin discrimination and cooperation in microbes. Annu Rev Microbiol 65:349–367. 10.1146/annurev.micro.112408.134109.

10. Lyons NA, Kolter R. 2017. *Bacillus subtilis* protects public goods by extending kin discrimination to closely related species. MBio 8:e00723–17. 10.1128/mBio.00723-17.

11. Kraigher B, Butulken M, Stefanic P, Mandic-Mulec I. 2021. Kin discrimination drives territorial exclusion during *Bacillus subtilis* swarming and restrains exploitation of surfactin. ISME J (in press). 10.1038/s41396-021-01124-4.

12. Lyons NA, Kraigher B, Stefanic P, Mandic-Mulec I, Kolter R. 2016. A combinatorial kin discrimination system in *Bacillus subtilis*. Curr Biol 26:733–742. 10.1016/j.cub.2016.01.032.

13. Mehdiabadi NJ, Jack CN, Farnham TT, Platt TG, Kalla SE, Shaulsky G, Queller DC, Strassmann JE. 2006. Kin preference in a social microbe. Nature 442:881–882. 10.1038/442881a.

14. Kalamara M, Spacapan M, Mandic-Mulec I, Stanley-Wall NR. 2018. Social behaviours by *Bacillus subtilis*: quorum sensing, kin discrimination and beyond. Mol Microbiol 110:863–878. 10.1111/mmi.14127.

15. Kearns DB, Losick R. 2004. Swarming motility in undomesticated *Bacillus subtilis*. Mol Microbiol 49:581–590. 10.1046/j.1365-2958.2003.03584.x.

16. Spacapan M, Danevcic T, Stefanic P, Porter M, Stanley-Wall NR, Mandic-Mulec I. 2020. The comX quorum sensing peptide of *Bacillus subtilis* affects biofilm formation negatively and sporulation positively. Microorganisms 8:1131. 10.3390/microorganisms8081131.

17. Mader U, Antelmann H, Buder T, Dahl MK, Hecker M, Homuth G. 2002. *Bacillus subtilis* functional genomics: genome-wide analysis of the DegS-DegU regulon by transcriptomics and proteomics. Mol Genet Genomics 268:455–467. 10.1007/s00438-002-0774-2.

18. Chowdhury SP, Hartmann A, Gao XW, Borriss R. 2015. Biocontrol mechanism by root-associated *Bacillus amyloliquefaciens* FZB42 - A review. Front Microbiol 6:780. 10.3389/fmicb.2015.00780.

19. Shao J, Xu Z, Zhang N, Shen Q, Zhang R. 2014. Contribution of indole-3-acetic acid in the plant growth promotion by the rhizospheric strain *Bacillus amyloliquefaciens* SQR9. Biol Fertil Soils 51:321–330. 10.1007/s00374-014-0978-8.

20. Xu Z, Zhang R, Wang D, Qiu M, Feng H, Zhang N, Shen Q. 2014. Enhanced control of cucumber wilt disease by *Bacillus amyloliquefaciens* SQR9 by altering the regulation of its DegU phosphorylation. Appl Environ Microbiol 80:2941–2950. 10.1128/AEM.03943-13.

21. Stefanic P, Belcijan K, Kraigher B, Kostanjsek R, Nesme J, Madsen JS, Kovac J, Sorensen SJ, Vos M, Mandic-Mulec I. 2021. Kin discrimination promotes horizontal gene transfer between unrelated strains in *Bacillus subtilis*. Nat Commun 12:3457. 10.1038/s41467-021-23685-w.

22. Wang D, Xu Z, Zhang G, Xia L, Dong X, Li Q, Liles MR, Shao J, Shen Q, Zhang R. 2019. A genomic island in a plant beneficial rhizobacterium encodes novel antimicrobial fatty acids and a self-protection shield to enhance its competition. Environ Microbiol 21:3455–3471. 10.1111/1462-2920.14683.

23. Chen XH, Vater J, Piel J, Franke P, Scholz R, Schneider K, Koumoutsi A, Hitzeroth G, Grammel N, Strittmatter AW, Gottschalk G, Süssmuth RD, Borriss R. 2006. Structural and functional characterization of three polyketide synthase gene clusters in *Bacillus amyloliquefaciens* FZB42. J Bacteriol 188:4024–4036. 10.1128/JB.00052-06.

24. Verhamme DT, Murray EJ, Stanley-Wall NR. 2009. DegU and Spo0A jointly control transcription of two loci required for complex colony development by *Bacillus subtilis*. J Bacteriol 911:100–108. 10.1128/JB.01236-08.

25. Xu Z, Xie J, Zhang H, Wang D, Shen Q, Zhang R. 2019. Enhanced control of plant wilt disease by a xylose-inducible *degQ* gene engineered into *Bacillus velezensis* strain SQR9XYQ. Phytopathology 109:36–43. 10.1094/PHYTO-02-18-0048-R.

26. Xu Z, Mandic-Mulec I, Zhang H, Liu Y, Sun X, Feng H, Xun W, Zhang N, Shen Q, Zhang R. 2019. Antibiotic bacillomycin D affects iron acquisition and biofilm formation in *Bacillus velezensis* through a Btr-mediated FeuABC-dependent pathway. Cell Rep 29:1192–1202. 10.1016/j.celrep.2019.09.061.

27. Xu Z, Shao J, Li B, Yan X, Shen Q, Zhang R. 2013. Contribution of bacillomycin D in *Bacillus amyloliquefaciens* SQR9 to antifungal activity and biofilm formation. Appl Environ Microbiol 79:808–815. 10.1128/AEM.02645-12.

28. Koumoutsi A, Chen XH, Vater J, Borriss R. 2007. DegU and YczE positively regulate the synthesis of bacillomycin D by *Bacillus amyloliquefaciens* strain FZB42. Appl Environ Microbiol 73:6953–6964. 10.1128/AEM.00565-07.

29. Lugtenberg B, Faina K. 2009. Plant-growth-promoting rhizobacteria. Annu Rev Microbiol 63:541–556. 10.1146/annurev.micro.62.081307.162918.

30. Santhanam R, Menezes RC, Grabe V, Li D, Baldwin IT, Groten K. 2019. A suite of complementary biocontrol traits allows a native consortium of root-associated bacteria to protect their host plant from a fungal sudden-wilt disease. Mol Ecol 28:1154–1169 10.1111/mec.15012.

31. Eng A, Borenstein E. 2019. Microbial community design: methods, applications, and opportunities. Curr Opin Biotechnol 58:117–128. 10.1016/j.copbio.2019.03.002.

32. Kong WT, Meldgin DR, Collins JJ, Lu T. 2018. Designing microbial consortia with defined social interactions. Nat Chem Biol 14:821–829. 10.1038/s41589-018-0091-7.

33. Riley MA, Wertz JE. 2002. Bacteriocins: evolution, ecology, and application. Annu Rev Microbiol 56:117–137. 10.1146/annurev.micro.56.012302.161024.

34. Kim PI, Ryu J, Kim YH, Chl YT. Production of biosurfactant lipopeptides iturin A, fengycin, and surfactin A from *Bacillus subtilis* CMB32 for control of *colletotrichum gloeosporioides*. J Microbiol Biotechn 20:138–145. 10.4014/jmb.0905.05007.

35. Fujita M, Losick R. 2005. Evidence that entry into sporulation in *Bacillus subtilis* is mediated by gradual activation of a master regulator spo0A. Genes Dev 19:2236–2244. 10.1101/gad.1335705.

36. Fujita M, González-Pastor JE, Losick R. 2005. High- and low-threshold genes in the Spo0A regulon of *Bacillus subtilis*. J Bacteriol 187:1357–1368. 10.1128/JB.187.4.1357-1368.2005.

37. Verhamme DT, Kiley TB, Stanley-Wall NR. 2007. DegU co-ordinates multicellular behaviour exhibited by *Bacillus subtilis*. Mol Microbiol 65:554–568. 10.1111/j.1365-2958.2007.05810.x.

38. Kobayashi K. 2007. Gradual activation of the response regulator DegU controls serial expression of genes for flagellum formation and biofilm formation in *Bacillus subtilis*. Mol Microbiol 66:395–409. 10.1111/j.1365-2958.2007.05923.x.

39. Wang P, Guo Q, Ma Y, Li S, Lu X, Zhang X, Ma P. 2015. DegQ regulates the production of fengycins and biofilm formation of the biocontrol agent *Bacillus subtilis* NCD-2. Microbiol Res 178:42–50. 10.1016/j.micres.2015.06.006.

40. Branda SS, González-Pastor JE, Ben-Yehuda S, Losick R, Kolter R. 2001. Fruiting body formation by *Bacillus subtilis*. Proc Natl Acad Sci U S A 98:11621–11626. 10.1073/pnas.191384198.

41. Landy M, Warren GH, Rosenmanm SB, Colio LG. 1948. Bacillomycin: An antibiotic from *Bacillus subtilis* active against pathogenic fungi. Proc Soc Exp Biol Med 67:539–541. 10.3181/00379727-67-16367.

42. Xu Z, Zhang H, Sun X, Lui Y, Yan W, Xun W, Shen Q, Zhang R. 2019. *Bacillus velezensis* Wall teichoic acids are required for biofilm formation and root colonization. Appl Environ Microbiol 85:e02116–02118. 10.1128/AEM.02116-18.

43. Zhou C, Shi L, Ye B, Feng H, Zhang J, Zhang R, Yan X. 2017. *pheS* *, an effective host-genotype-independent counter-selectable marker for marker-free chromosome deletion in *Bacillus amyloliquefaciens*. Appl Microbiol Biotechnol 101:217–227. 10.1007/s00253-016-7906-9.

44. Hamon MA, Lazazzera BA. 2001. The sporulation transcription factor Spo0A is required for biofilm development in *Bacillus subtilis*. Mol Microbiol 42:1199–1209. 10.1046/j.1365-2958.2001.02709.x.

45. Nicholson WL. 2008. The *Bacillus subtilis ydjL* (*bdhA*) gene encodes acetoin reductase/2,3-butanediol dehydrogenase. Appl Environ Microbiol 74:6832–6838. 10.1128/AEM.00881-08.

46. Li Q, Li Z, Li X, Xia L, Zhou X, Xu Z, Shao J, Shen Q, Zhang R. 2018. FtsEX-CwlO regulates biofilm formation by a plant-beneficial rhizobacterium *Bacillus velezensis* SQR9. Res Microbiol 169:166–176. 10.1016/j.resmic.2018.01.004.

47. Fan B, Chen XH, Budiharjo A, Bleiss W, Vater J, Borriss R. 2011. Efficient colonization of plant roots by the plant growth promoting bacterium *Bacillus amyloliquefaciens* FZB42, engineered to express green fluorescent protein. J Biotechnol 151:303–311. 10.1016/j.jbiotec.2010.12.022.

